# Age-specific Aβ-Tau interactions underlie spatial memory deficits in an APP/PS1 amyloidosis model

**DOI:** 10.64898/2025.12.06.689974

**Authors:** Athanasios Metaxas, Marco Anzalone, Ramanan Vaitheeswaran, Vivien Tejsi, Shuoshuo Li, Georgia Mitilineou, Kate L. Lambertsen, Hélène Audrain, David J. Brooks, Anne M. Landau, Bente Finsen

## Abstract

**Background:** Amyloid-β (Aβ) and tau pathology are key molecular hallmarks of Alzheimer’s disease (AD), yet how their interaction contributes to cognitive decline remains unclear. We investigated the relationship between Aβ burden, tau pathology, presynaptic density, and spatial learning and memory in the *APP*_swe_/*PSEN1*_ΔE9_ (APP/PS1) transgenic (TG) mouse model of amyloidosis.

**Methods:** Spatial learning and memory were assessed with the Barnes maze test in male TG and wild-type (WT) littermate mice, aged 6, 12, and 18 months. Gross visual function was assessed indirectly in 18-month-old animals using the light/dark exploration test. Brains were collected for autoradiography of tau pathology and presynaptic density with [^18^F]Flortaucipir and [^3^H]UCB-J, respectively, while Aβ plaque load was measured by immunohistochemistry. Correlation and linear mixed-effects regression analyses were used to assess relationships between behavioral and pathological measures.

**Results:** APP/PS1 mice showed normal cognitive performance at 6 months, a selective long-term memory deficit at 12 months, and severe impairments in learning and retention at 18 months, independent of visual confounds. Age-dependent increases in [^18^F]Flortaucipir binding and Aβ plaque load were observed in all brain regions of TG compared to WT mice, whereas [^3^H]UCB-J binding was increased in a region-dependent manner in 18-month-old TG vs. WT animals. Barnes maze performance during the final day of testing correlated negatively with both Aβ and tau pathology across all areas examined. Linear regression revealed a significant association between tau and age and between tau and Aβ in the cortex, indicating that memory decline in ageing TG mice was driven by the combined effects of these pathologies.

**Conclusions:** Deficits in memory retention precede impairments in task learning performance in APP/PS1 mice. Spontaneous tau accumulation contributes to the progressive cognitive decline, capturing key aspects of the Aβ-tau interaction observed in human AD.

## Introduction

Alzheimer’s disease (AD) is a neurodegenerative disorder that is characterized by the extracellular deposition of amyloid-β (Aβ) peptides into plaques, the intracellular accumulation of hyperphosphorylated tau protein into neurofibrillary tangles (NFTs), a brain-specific form of inflammation, and widespread synaptic dysfunction and neuronal loss, all of which contribute to the progressive cognitive decline observed in AD patients [1]. Among the pathological hallmarks of AD, synaptic loss is considered to be the strongest structural correlate of cognitive impairment, possibly reflecting the cumulative impact of Aβ and tau pathology on neuronal connectivity [2, 3].

Neuroimaging studies show that the accumulation of Aβ is an early pathogenic event in AD, whereas cortical tau pathology, particularly the formation of NFTs, typically emerges later in the disease course, correlating to a larger extent with cognitive decline than Aβ [4, 5]. Mounting evidence suggests that the accumulation of Aβ and the initiation and spread of tau pathology are mechanistically linked in AD. For example, preclinical studies have shown that Aβ pathology can enhance tau phosphorylation, aggregation and propagation, ultimately disrupting synaptic function [6–8]. Recent clinical trials of anti-amyloid monoclonal antibodies, including aducanumab, lecanemab, and donanemab, have further demonstrated reduced levels of phosphorylated tau in the CSF and plasma of treated patients, slower accumulation of tau pathology on positron emission tomography (PET) imaging, and a correlation between amyloid clearance, tau biomarker improvements, and clinical benefit [9]. Together, these findings support a model in which Aβ-driven tau pathology mediates synaptic degeneration and cognitive decline in AD.

Mouse models of familial AD recapitulate many aspects of amyloid pathology and have been extensively used to study Aβ-induced synaptic dysfunction and neuroinflammation [10]. Although these models do not typically exhibit extensive tau pathology, we have previously shown that they can develop tau-related alterations characteristic of AD under certain conditions. For instance, *APP*_swe_/*PSEN1*_de9_ (APP/PS1) mice spontaneously develop aggregates of hyperphosphorylated tau in the neocortex and hippocampus at the age of 18-24 months [11], when their cerebral amyloid burden is highest [12]. Moreover, in a recent systematic review and meta-analysis we have shown that tau phosphorylation in amyloidosis models is a dynamic, region-specific process that is closely associated with the accumulation of Aβ, and, depending on the mouse model studied, can resemble either the early or later stages of tau pathology observed in human AD [13]. These data suggest that Aβ deposition may be sufficient to trigger tau misprocessing and aggregation, even in the absence of tau overexpression, rendering amyloidosis models appropriate for studying the interplay between Aβ pathology, tau alterations, synaptic integrity and cognitive decline in AD.

In the present study, we sought to investigate whether impairments in the Barnes maze test of spatial learning and memory [14] correlate with molecular markers of tau pathology, synaptic integrity and amyloid burden in APP/PS1 mice. Defining how hallmark AD pathologies contribute to cognitive impairment in an amyloidosis model is crucial for identifying and mitigating the mechanisms most closely linked to functional decline in AD, and for enhancing the model’s translational relevance.

## Materials and Methods

### Animals and Ethics

All procedures complied with Danish law (Dyreværnsloven-Protection of Animals Act, nr 344/2005) and European Union directive 2010/63/EU, regulating animal research. Ethical permission was granted by the Animal Ethics Inspectorate of Denmark (nr. 2016/15-0201-00952).

APP/PS1 transgenic (TG) mice, originally generated by Jankowsky et al. 2001 [15], were bred and maintained as hemizygotes on a C57BL/6J background in the Biomedical Laboratory of the University of Southern Denmark. Age- and sex-matched wild-type (WT) littermate mice were used as controls. The animals were group-housed (8 per cage) under controlled temperature (21±1°) and humidity (45-65%) conditions, and a 12:12 h light/dark cycle (lights on: 7 am). Food and water were provided *ad libitum*. All experiments were conducted during the light phase of the cycle, between 08:00 h and 15:00 h.

Male TG and WT mice, aged 6, 12 and 18 months were used in the studies (n=9-11/genotype and age-group for the Barnes maze assay; n=6/group for the light aversion test, total number of animals used: 70). These time points were chosen to reflect the early, established and advanced phases of cerebral amyloidosis in the APP/PS1 TG mouse model [12]. At least five days prior to behavioral testing, mice were transferred to a dedicated behavioral testing facility and habituated to handling daily to minimize stress during subsequent assessments.

### Barnes maze test

The Barnes maze test is a dry-land alternative to the Morris water maze test and was used to assess spatial learning and memory, as previously described [16]. The Barnes maze apparatus (Panlab S.L.U., Barcelona, Spain) was a circular platform with 20 equidistant holes around its perimeter, elevated 1.5 m above the floor and brightly illuminated by four 120W halogen lamps, to create an aversive environment. One of the 20 peripheral holes led to a recessed, dark escape box (goal box), providing animals with a route to escape from the brightly lit arena.

On day 0, animals were habituated to the Barnes maze arena. Mice were placed in a transparent tube at the centre of the maze for 30 s, guided to the escape hole within 10-15 s and allowed 2 min to climb down and enter the dark goal box. If they failed to enter, mice were gently pushed inside the goal box using the transparent tube and remained there for 2 min before being returned to their home cage. Spatial learning was assessed over three consecutive days (days 1–3; acquisition phase), with each mouse receiving five training trials per day, each lasting 180 s. For starting point randomization, each mouse was placed for 15 s at the centre of the maze, within an opaque starting cylinder. The lights were then turned on, the cylinder lifted, and the animal was allowed 180 s to locate the escape box. If successful, the mouse remained in the goal box for 1 min before being returned to its home cage; if not, it was gently guided into the box and kept there for 1 min before being returned to its home cage. Between trials, the maze and escape box were cleaned and dried thoroughly, by rinsing with 70% ethanol and water. Measurements during the acquisition phase included total latency, which was the time required for an animal to enter the escape box (all 4 paws inside the box), and primary latency, which was the time required for an animal to reach the escape hole for the first time (first correct nose-poke).

Spatial memory was assessed 24 h (day 4) and 8 days after the acquisition phase (day 11), using single-probe trials at each time-point. The escape box was removed for these sessions. Mice were placed in opaque starting cylinder for 15 s, the lights turned on, and the animals allowed 90 s to explore the maze. In addition to primary latency, measurements during the memory-probing phase included target block time, which was the time each mouse spent in the quarter of the arena where the escape box used to be situated, nose-poke distribution, which was the number of nose-pokes conducted by each animal in each of the 20 holes during the 90 s trial, and primary errors, which was the number of incorrect holes a mouse visited before reaching the target (escape) hole for the first time.

For the Barnes maze experiments, mice were always tested in a paired manner, with one TG and one WT mouse completing a trial before proceeding to the next pair. Where possible, testing followed an age-paired design (e.g., TG and WT mice at 6, 12, and 18 months tested in alternating sequence). Data were generated from eight independent Barnes maze experiments, all performed under blinded conditions and recorded using a digital video camera. The location of the escape box was kept constant during a given experiment but was changed to different positions between experiments, to control for fixed environmental cues (location bias). Data were analysed in a blinded manner by two independent experimenters, and mean values from their observations were used for subsequent analysis.

### Light aversion test

The light aversion test was performed according to Thomson et al. 2010 [17], using a separate cohort of male, 18-month-old WT and APP/PS1 mice (n=6/group). The test was used to provide an indirect assessment of gross visual function, by confirming that mice can detect and respond to bright, aversive visual stimuli. The animals were tested in a 40 x 40 x 40 cm arena, which was divided into two equally sized parts, connected at the mid-point of a dividing, non-transparent wall by a small 7 x 7 cm portal. An open-top chamber comprised the brightly lit compartment, which was illuminated using identical conditions as in the Barnes maze assay. The top of the dark compartment was constructed by transparent polypropylene material with a red tint, allowing the experimenter to observe behavior in the dark chamber.

Animals were tested in the light/dark assay in a paired and counterbalanced manner. The test began by gently placing a mouse in the middle of either the dark or light compartment, allowing it to freely explore the arena for a period of 10 min. Time spent in a specific compartment was measured as an index of light aversion. All four mouse paws had to enter the dark or illuminated chamber for an animal to be considered as being inside a specific compartment. The chamber was cleaned in between sessions with 70% ethanol and water. Animal behavior was recorded and analysed by two independent observers, one of which was blinded as to group genotype. A mean value of time spent in each compartment was calculated from the observations of the two experimenters and later used for analysis.

### Tissue sectioning

Immediately following the memory probing session on Barnes maze Day 11, the mice were euthanized by cervical dislocation, and the brains immediately removed and frozen in isopentane on dry-ice (−30°C). Tissue sectioning was carried out at −17°C in a Leica CM3050S cryostat (Nussloch, Germany). Brains were mounted in the coronal plane and a series of 20 μm-thick sections were collected from forebrain to hindbrain levels, at 300 μm intervals. The sections were mounted onto Superfrost^TM^ Plus microscope slides (ThermoScientific, Denmark), dried at 4°C in a box containing silica gel for at least 2 h, and stored at −80°C for subsequent autoradiography and immunohistochemistry experiments.

### Quantification of tau pathology

These experiments were conducted at the Department of Nuclear Medicine and PET-centre, University of Aarhus, Denmark. The paired-helical-filament tau tracer [^18^F]Flortaucipir was synthesized according to established methods [18], and used as described in Metaxas et al. 2019 [11]. Sections from APP/PS1 mice and aged-matched WT animals were thawed to room temperature (RT) for 20 min and fixed/permeabilised in 100% methanol for 20 min. The sections were incubated for a period of 60 min in a 160 mL bath of 10 mM phosphate buffered saline (PBS, pH 7.4), containing 443.0±119.3 kBq/mL [^18^F]Flortaucipir (specific activity: 144±46 GBq/μmol). A series of adjacent sections was incubated with identical amounts of radioligand in the presence of 15 μM ‘cold’ flortaucipir, to assess non-specific binding (NSB). Following incubation, sections were serially rinsed in PBS (1 min), 70% ethanol in PBS (2 x 1 min), 30% ethanol in PBS (1 min) and PBS (1 min). After rapid drying under a stream of cold air, the sections were placed in light-tight cassettes and exposed against FUJI multi-sensitive phosphor screens for 30 min (BAS-IP SR2025, GE Healthcare Life Sciences). To allow quantification, standards of known radioactive concentration were prepared by serial dilution of the [^18^F]Flortaucipir incubation solution and exposed along with the sections. Images were developed using a BAS-5000 phosphor-imager at 25 μm resolution. Values of specific binding were derived after subtraction of non-specific from total binding images.

### Quantification of synaptic density

[^3^H]UCB-J was used to quantify synaptic vesicle glycoprotein 2A (SV2A), a marker of presynaptic density in the brains of WT and TG mice, as described in Metaxas et al. 2019 [19]. Sections were thawed to RT and pre rinsed in 50 mM Tris–HCl buffer (pH 7.4), containing 150 mM NaCl, 5 mM KCl, 1.5 mM MgCl2, and 1.5 mM CaCl2 (2 x 10 min). The sections were then incubated for 2 h in assay buffer, containing 1 nM [^3^H]UCB-J (specific activity 82.0 Ci/mmol; NT1099, NOVANDI Chemistry AB). NSB was determined in the presence of excess levetiracetam (TOCRIS), which was used at the concentration of 500 μM [i.e., ∼ 500 x levetiracetam’s IC_50_ for SV2A as determined in [^3^H]ucb 30889 competition binding [20]]. Incubations were terminated by three 1-min rinses in ice-cold 50 mM Tris-HCl buffer (pH 7.4), followed by a rapid rinse in ice-cold dH_2_O. The sections were rapidly dried and placed on Carestream® KodakR BioMax MR film for 4 weeks. To allow quantification, ^3^H microscales of known radioactive concentration were also exposed to film (American Radiolabeled Chemicals, Inc). The films were developed with substitute KODAK D-19 developer (TED PELLA, Inc), washed in dH_2_O, and fixed in Carestream® autoradiography GBX fixer. Images were digitized using a white sample tray and the Coomassie Blue settings on a ChemiDoc^TM^ MP imaging system (BIO-RAD) and analysed in ImageJ (version 1.51; National Institutes of Health, MD, United States). Mean grey values were extracted from manually drawn regions of interest following background subtraction and converted to fmol/mg tissue equivalent using the co-exposed calibration standards.

### Quantification of amyloid pathology

Preliminary autoradiography for Aβ plaques was conducted using the amyloid-preferring Pittsburgh compound B (PIB), which was radiolabelled with ^11^C at the Department of Nuclear Medicine and PET-centre, University of Aarhus, Denmark, as previously described [21]. In line with previous reports of limited tracer sensitivity in APP/PS1 mice [22, 23], no specific signal was detected in these preliminary studies (Supplementary Fig. 1). Therefore, Aβ burden was quantified by immunohistochemistry with the 6E10 antibody (SIG-39340, Nordic BioSite, Denmark). The 6E10 clone is raised against amino acids 1-16 of human Aβ and recognizes multiple amyloid peptides as well as precursor forms.

Fresh-frozen sections were fixed overnight at 4°C in 10% neutral buffered formalin (NBF). They were then immersed in 70% formic acid for 30 min, rinsed once for 10 min in 50 mM Tris-buffered saline (TBS), and further washed in TBS containing 1% Triton X-100 (TBS-Tx, 3 × 15 min). After incubation for 30 min in TBS-Tx supplemented with 10% fetal bovine serum (FBS), the sections were incubated overnight at 4°C with the biotinylated 6E10 antibody. The primary antibody was diluted 1:500 in TBS containing 10% FBS. On the following day, sections were brought to RT for 30 min and washed in TBS-Tx (3 × 15 min). Endogenous peroxidase activity was subsequently quenched using a TBS/methanol/H₂O₂ solution (8:1:1) for 20 min. After washing in TBS-Tx (3 × 15 min), sections were incubated for 3 h with horseradish peroxidase (HRP)-conjugated streptavidin (1:200; GE Healthcare Life Sciences, Denmark) in TBS + 10% FBS. Following a final rinse in TBS (3 × 10 min), staining was developed using 0.05% 3,3′-diaminobenzidine (DAB) in TBS containing 0.01% H₂O₂2 (Sigma Aldrich Co., Denmark). Sections were then thoroughly rinsed in distilled water, dehydrated through a graded ethanol series (70%, 96%, 99%), cleared in xylene, and cover slipped using PERTEX (HistoLab, Denmark).

Digital images were obtained under the 4x objective of an Olympus DP80 Dual Monochrome CCD camera, mounted on a motorized BX63 Olympus microscope. For analysis, the images were converted to 8-bit and manually thresholded in ImageJ. The particle analysis plugin was used to measure the percentage of immunoreactive area relative to total image area (% area plaque load).

### Data analysis & statistics

Total latency, primary latency, time spent in the target block, primary errors and nose-poke distribution were analyzed during the acquisition and/or memory-probing phases of the Barnes maze with a mixed-model ANOVA, with age and genotype as the between-subject variables, and time (day) as the within-subject, repeated variable. For the light aversion assay, time spent in the illuminated and dark compartments was analyzed using a mixed-model ANOVA, with chamber (dark vs. illuminated) as the within-subject variable and genotype as the between-subject variable. For autoradiographic and immunohistochemistry measures, factorial ANOVAs were conducted as appropriate, with genotype, age, and brain region as the independent factors. *Post hoc* comparisons were performed using the least significant difference (LSD) test when main or interaction effects reached significance.

Correlation analyses between behavioral parameters on day 11 and [^18^F]Flortaucipir binding, [^3^H]UCB-J binding and Aβ burden were performed using Pearson’s correlation coefficient. To account for age as a potential confounding factor, we subsequently applied linear mixed-effects regression models, including age, Αβ, tau and synaptic measures as fixed predictors and animal identity as a random effect. This approach allowed us to determine whether variability in pathology measures predicted cognitive performance independently of ageing. Cortical regions (frontal, parietal/temporal, and occipital) were analyzed together, and the hippocampus was assessed separately. In all cases, statistical significance was set at *P* < 0.05, and all analyses were carried out using SPSS (IBM) or Statistica (v12; StatSoft Inc). Data are presented as mean ± SEM of 9-11 animals/group.

## Results

### Learning impairment in ageing APP/PS1 mice

Primary latency, which is the time required for a mouse to reach the escape hole for the first time, was measured in groups of 6-, 12-, and 18-month-old WT and TG animals on days 1-3, i.e., throughout the acquisition phase of the Barnes maze test (Fig. 1). There was no difference in primary latency between WT and APP/PS1 mice at 6 and 12 months of age. At 18 months, TG mice had increased primary latency over age-matched WT controls on training days 1 and 2 (*P*<0.01; LSD *post-hoc* tests). Moreover, on all training days, 18-month-old TG mice took longer to reach the escape hole compared to 6-month-old APP/PS1 mice (LSD *post-hoc* tests). No effect of age on the learning behavior of WT animals was observed. Repeated measures ANOVA confirmed significant effects of age [F(2,51)=5.3; *P*<0.01], genotype [F(1,51)=8.8; *P*<0.01] and training day [F(2,102)=50.7; *P*<0.001] on the primary latency of WT and TG animals, as well as significant genotype x age interaction effects [F(2,51)=3.7; *P*<0.05]. Similarly, analysis of total latency, defined as the time required for a mouse to enter the escape box, revealed significant main effects of age [F(2,49)=17.8; *P*<0.001], genotype [F(1,49)=8.8; *P*<0.01] and training day [F(2,98)=194.9; *P*<0.001], as well as significant genotype x age interaction effects [F(2,49)=3.6; *P*<0.05] (Supplementary Fig. 2).

**Figure 1.**
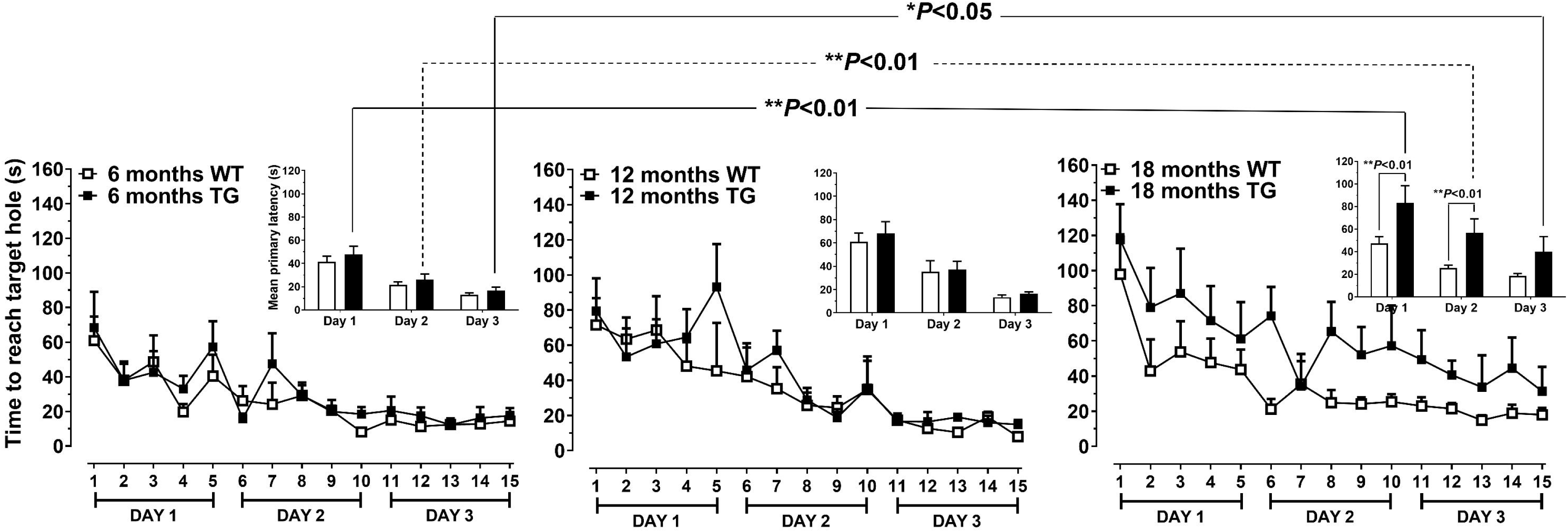
Learning impairment in ageing APP/PS1 mice. Primary latency, defined as the time required for a mouse to reach the target hole for the first time, was measured in wild-type (WT, open squares) and APP/PS1 transgenic (TG, filled squares) mice during three consecutive training days (five trials per day) in the Barnes maze. At 6 and 12 months of age, no significant differences in primary latency were observed between WT and TG mice, indicating preserved learning performance. At 18 months, TG mice exhibited significantly longer latencies compared to age-matched WT controls on training days 1 and 2 (*P*<0.01; LSD post hoc test). The insets show mean primary latency (±SEM) for each training day across ages, with 18-month-old TG mice performing worse than 6-month-old animals. Repeated-measures ANOVA confirmed significant main effects of genotype [F(1,51)=4.7; *P*<0.05] and significant trial day x age interaction effects on the primary latency of TG and WT animals [F(2,51)=7.5; *P*<0.01]. Data are expressed as mean ± SEM (n = 9-11 per group).

### Deficits in memory retention in APP/PS1 mice

Spatial memory retention was assessed 24 h and 8 days after the end of the acquisition phase, on days 4 and 11 after experiment initiation, respectively. Genotype- and age-specific deficits in the behavior of WT and TG animals were observed (Fig. 2A-C).

**Figure 2.**
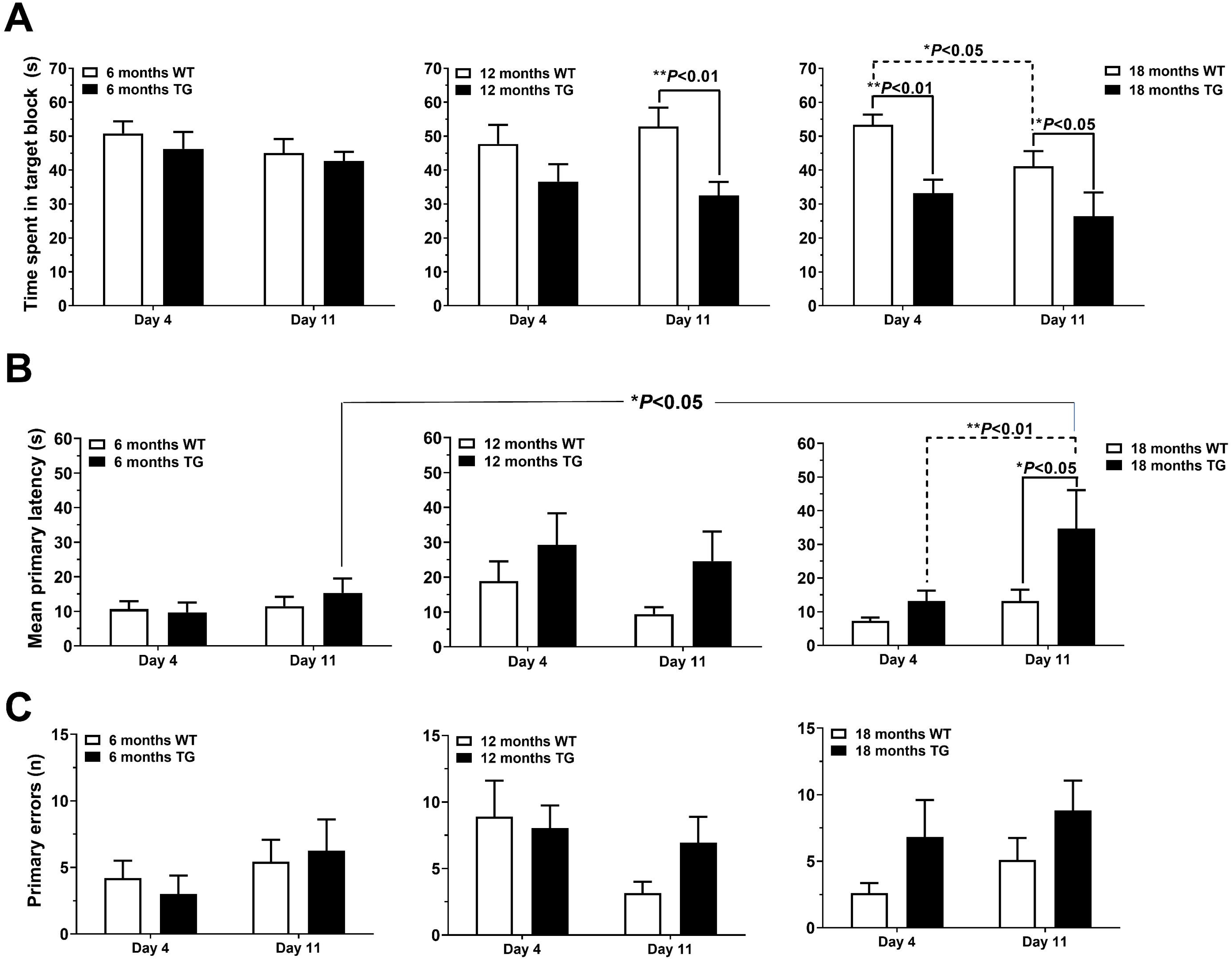
Deficits in memory retention in ageing APP/PS1 mice. Spatial memory performance was probed in wild-type (WT, open bars) and APP/PS1 transgenic (TG, filled bars) mice, 24 hours (Day 4) and 8 days (Day 11) after the end of training. (A) Time spent in the target block (the maze quadrant containing the escape hole during training) was reduced in 12-month-old TG mice compared to WT on Day 11 (*P*<0.01), and in 18-month-old TG vs. WT mice on both Day 4 (*P*<0.01) and Day 11 (*P*<0.05). Target-block time also decreased between Days 4 and 11 in 18-month-old WT mice (*P*<0.05). (B) Mean primary latency, defined as the time to reach the target hole for the first time, was increased in 18-month-old TG mice compared to WT on Day 11 (*P*<0.05), compared to 6-month-old TG mice on Day 11 (*P*<0.05) and between Day 4 and Day 11 (*P*<0.001), indicating progressive impairment in long-term memory retention. (C) Primary errors, defined as the number of incorrect holes visited before reaching the target hole, did not differ between genotypes at any age or time point. Data represent mean ± SEM of n = 9-11 animals per group.

Target block time, which was the time each mouse spent in the quarter of the arena where the escape box used to be situated, was reduced in 12-month-old TG vs. WT mice on trial day 11 (*P*<0.01; LSD *post-hoc* tests; Fig. 2A). Eighteen-month-old TG mice spent less time in the target block compared to WT animals on both day 4 (*P*<0.01) and day 11 (*P*<0.05; LSD *post-hoc* tests). Residence time in the target block was also reduced in 18-month-old WT mice on day 11 vs. day 4 (*P*<0.05; LSD *post-hoc* tests). The overall repeated measures ANOVA showed significant main effects of genotype on target-block time [F(1,51)=16.3; *P*<0.001], with tendencies for significant age [F(2,51)=2.1; *P*=0.13], trial day [F(1,51)=3.6; *P*=0.06] and genotype x age interaction effects [F(2,51)=2.1; *P*=0.13].

For primary latency (Fig. 2B), 18-month-old TG mice showed greater latency times compared to age-matched WT animals on day 11 (*P*<0.05) and took longer to reach the escape hole on day 11 vs. day 4 (*P*<0.001, LSD *post-hoc* tests). In addition, primary latency was increased in 18- vs. 6-month-old TG mice on day 11 (*P*<0.05; LSD *post-hoc* tests). The overall ANOVA confirmed significant main effects of genotype [F(1,51)=4.7; *P*<0.05] and significant trial day x age interaction effects on the primary latency of TG and WT animals [F(2,51)=7.5; *P*<0.01]. The genotype x trial day interaction effect was nearly significant [F(1,51)=3.5; *P*=0.06]. The number of primary errors (Fig. 2C) was not different between TG and WT animals [genotype effect: F(1,50)=2.1; *P*=0.15], on any of the days [genotype x day interaction effect: F(1,50)=1.2; *P*=0.28] and ages examined [genotype x age interaction effect: F(2,50)=1.0; *P*=0.38].

Progressive deficits in the performance of both WT and TG animals were further evident when the distribution of nose-pokes across the holes of the Barnes maze platform was analyzed on trial days 4 and 11 (Fig. 3A-C). Although the total number of nose-pokes conducted during the memory-probing sessions was not different between WT and TG animals (Fig. 3A-C, inserts), their distribution differed significantly between genotypes, starting at 12 months of age. Thus, 12-month-old APP/PS1 animals made fewer nose-pokes in the target hole compared to age-matched WT controls on day 11 (Fig. 3B). In 18-month-old mice, differences in the number of targeted nose-pokes between WT and TG mice were observed on day 4. Moreover, 18-month-old WT animals performed less nose-pokes in the target hole on day 11 compared to day 4 (Fig. 3C). Repeated measures ANOVA confirmed significant main effects of genotype [F(1,102)=8.4; *P*<0.01] and age [F(2,102)=6.6; *P*<0.01] on the distribution of nose-pokes, along with significant hole x genotype [F(19,1938)=7.2; *P*<0.001], hole x age [F(38,1938)=3.1; *P*<0.001] and hole x trial day interaction effects [F(19,1938)=3.3; *P*<0.001].

**Figure 3.**
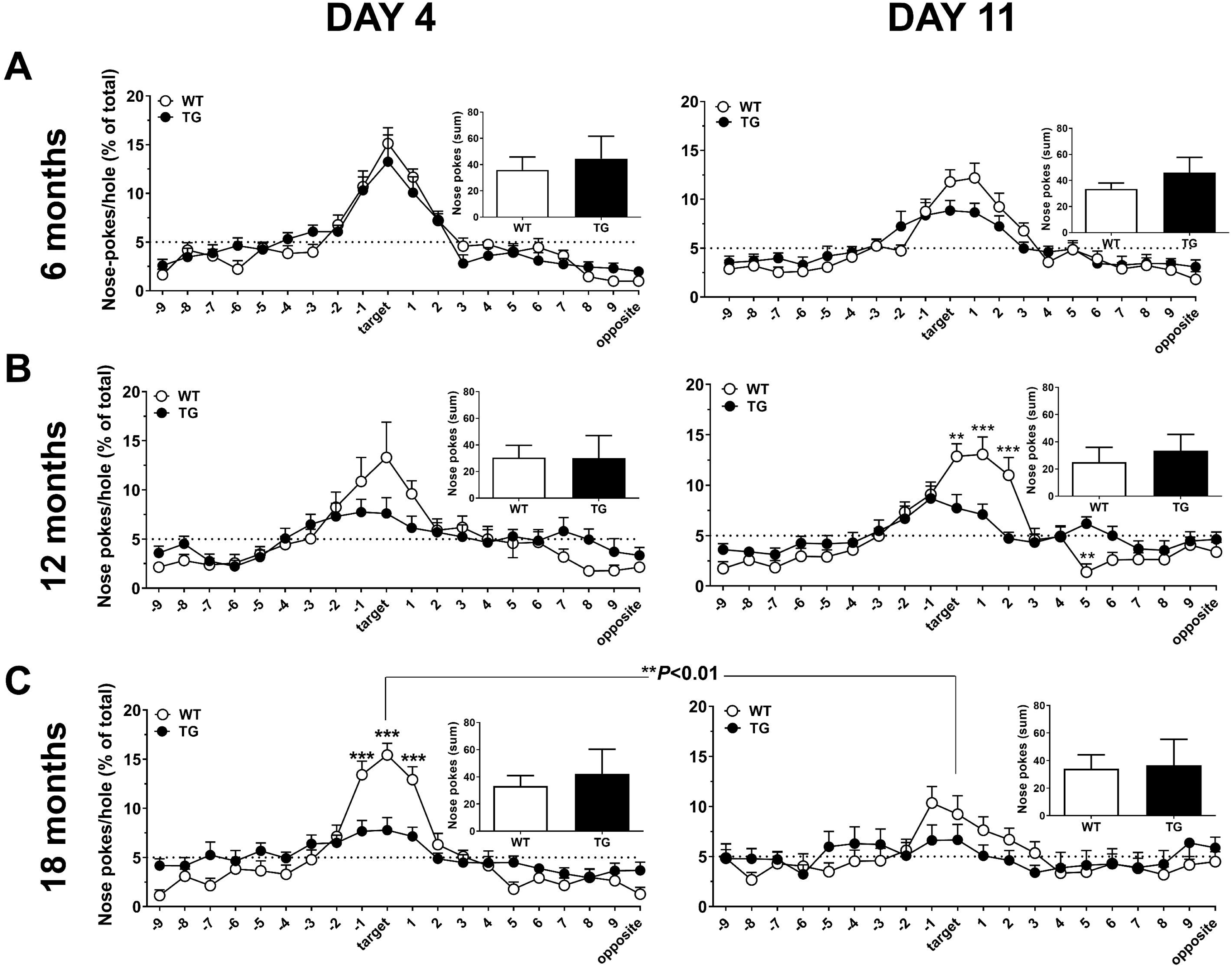
Distribution of nose-pokes during Barnes maze probe trials. The spatial distribution of nose-pokes across all holes of the Barnes maze was analyzed in wild-type (WT, open circles) and APP/PS1 transgenic (TG, filled circles) mice during the probe trials, which were conducted 24 hours (Day 4) and 8 days (Day 11) after training. The dashed horizontal line represents the level of nose-poking expected by chance (random performance). Insets show the total number of nose-pokes performed during each session. (A) At 6 months of age, WT and TG mice showed comparable search patterns and similar total nose-poke counts. (B) At 12 months, TG mice made significantly fewer nose-pokes in the target hole compared to WT animals on Day 11 (*P*<0.01–0.001), indicating impaired memory for the target location. (C) At 18 months, TG mice exhibited impaired spatial memory, as shown by reduced preference for the target hole compared to WT mice on Day 4 (*P*<0.001). WT animals also showed reduced target-directed responding between Days 4 and 11 (*P*<0.01). Repeated measures ANOVA confirmed significant main effects of genotype [F(1,102)=8.4; *P*<0.01] and age [F(2,102)=6.6; *P*<0.01] on the distribution of nose-pokes, along with significant hole x genotype [F(19,1938)=7.2; *P*<0.001], hole x age [F(38,1938)=3.1; *P*<0.001] and hole x trial day interaction effects [F(19,1938)=3.3; *P*<0.001]. Data are expressed as mean ± SEM of n = 9-11 mice per group.

### Lack of genotype-induced effects in the light aversion test

To exclude the possibility that visual impairment contributed to the cognitive deficits observed in the Barnes maze arena, independent groups of 18-month-old WT and TG animals were tested in the light aversion assay. Both WT (*P*<0.05) and TG mice (*P*<0.01; LSD *post-hoc* tests) displayed a strong preference for the dark relative to the illuminated compartment of the arena, with no differences observed between genotypes (Fig. 4). Repeated measures ANOVA confirmed a significant main effect of compartment [F(1,10)=8.7, *P*=0.01], but no significant genotype [F(1,10)=1.0, *P*=0.34] or genotype × compartment interaction effects [F(1,10)=0.0, *P*=0.84] on the time-spent in the light vs. dark area of the arena.

**Figure 4.**
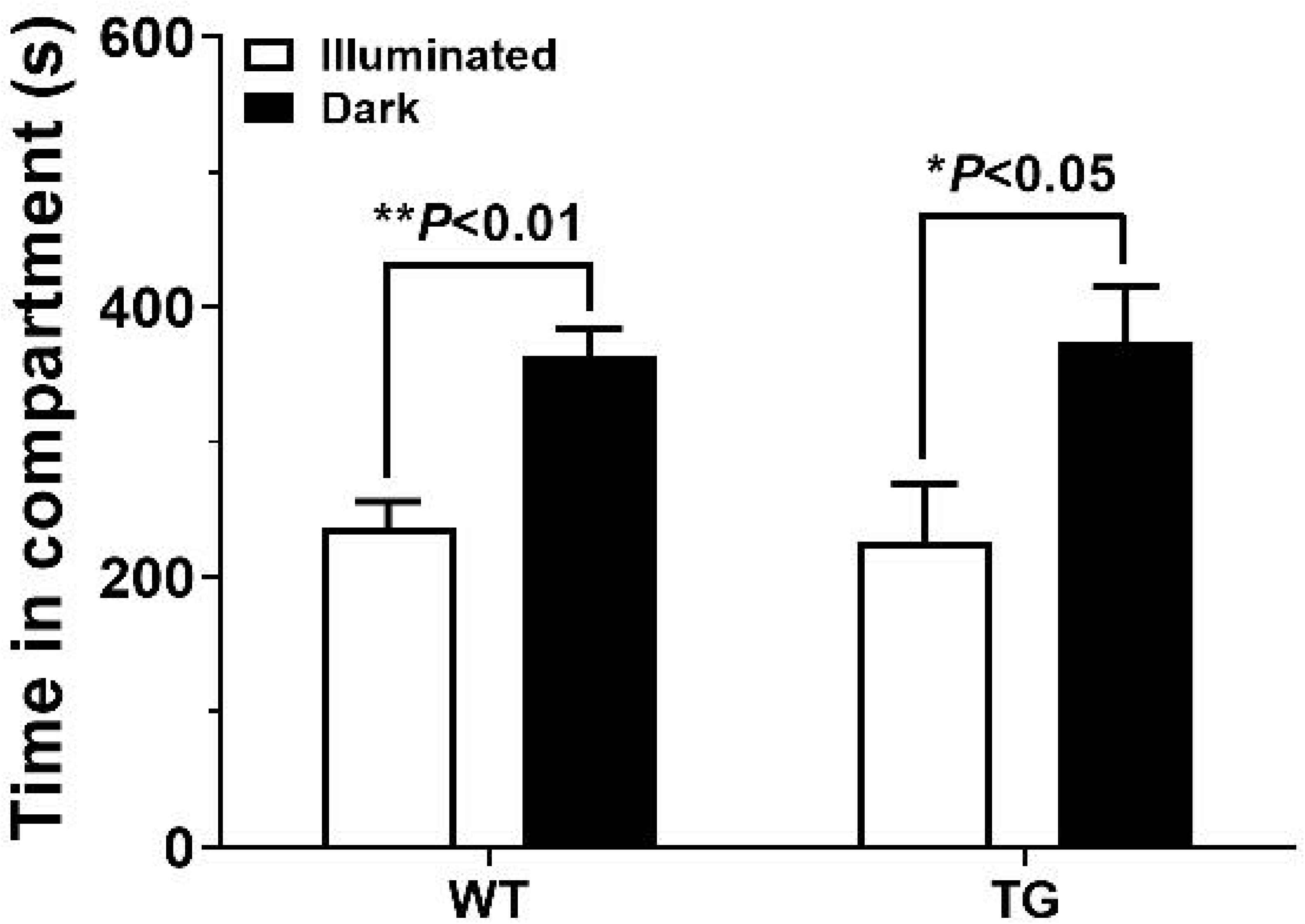
Lack of visual impairment in the light/dark exploration test. Time spent in the illuminated (open bars) and dark (filled bars) compartments was measured in 18-month-old wild-type (WT) and APP/PS1 transgenic (TG) mice. Both WT (*P*<0.01) and TG (*P*<0.05) mice spent significantly more time in the dark compartment. No genotype-induced differences were observed between genotypes, suggesting that visual function were comparable across groups. Data are presented as mean ± SEM of n = 6 mice per group.

### Age-dependent increase in [^18^F]Flortaucipir binding in TG mice

Immediately following the Barnes maze probe trial on day 11, mice were euthanized by cervical dislocation, and their brains isolated and processed for autoradiography with the tau tracer [^18^F]Flortaucipir. Progressive, age-dependent increases in the binding levels of [^18^F]Flortaucipir were observed in APP/PS1 mice exclusively, across all brain areas quantified (Fig. 5A). In 18-month-old TG mice, the highest levels of specific binding were observed in the frontal cortex, and the lowest in the hippocampus. No age-related changes were observed in WT mice. Levels of [^18^F]Flortaucipir binding were significantly higher in TG compared to age-matched WT mice at 12 and 18 months of age (*P*<0.001; LSD *post-hoc* tests). Factorial ANOVA revealed significant main effects of age [F(2,204)=163.4, *P*<0.001], genotype [F(1,204)=656.2, *P*<0.001] and brain region [F(3,204)=30.8, *P*<0.001] on the binding levels of [^18^F]Flortaucipir, along with significant age x genotype interaction effects [F(2,204)=128.9, *P*<0.001]. Representative autoradiograms of [^18^F]Flortaucipir binding sites are shown in Fig. 5B.

**Figure 5.**
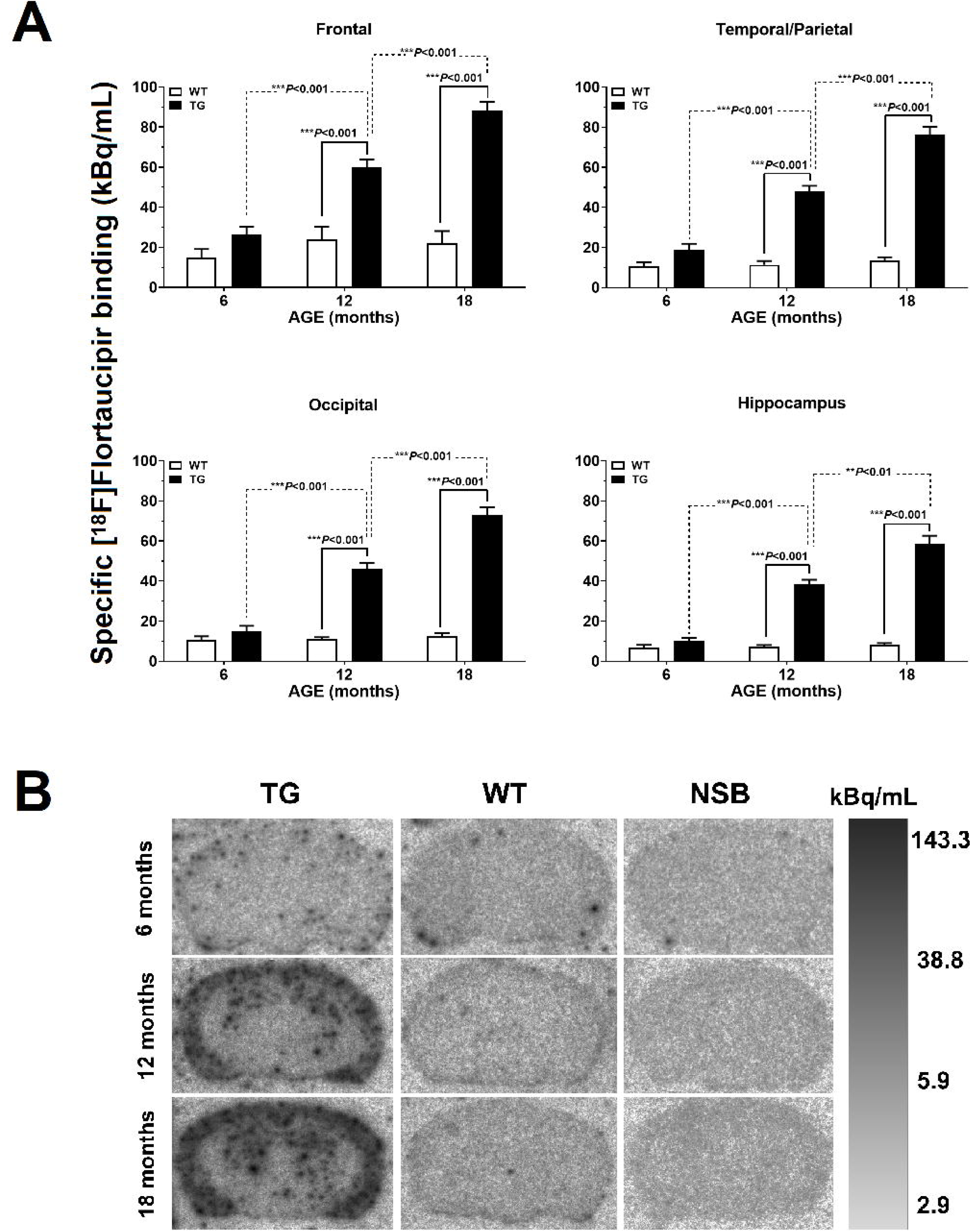
Age-dependent increase in [^18^F]Flortaucipir binding in APP/PS1 mice. (A) Specific binding of the tau tracer [^18^F]Flortaucipir was quantified in the frontal, temporal/parietal, occipital cortices, and hippocampus of wild-type (WT, open bars) and APP/PS1 transgenic (TG, filled bars) mice at 6, 12, and 18 months of age. A robust, age-dependent increase in [^18^F]Flortaucipir binding was observed in TG mice across all regions examined (*P*<0.001), with significantly higher binding levels in TG compared with WT animals at 12 and 18 months of age (*P*<0.001). No age-related changes were detected in WT mice. (B) Representative autoradiograms show regional increases of [^18^F]Flortaucipir binding in 12- and 18-month-old TG vs. age-matched WT mice. Non-specific binding (NSB) sections were incubated in the presence of excess unlabelled ligand. The scale bar shows a grayscale representation of binding intensity, expressed in kBq/mL.

### Age-dependent increase in [^3^H]UCB-J binding in TG mice

Preynaptic density was assessed using [^3^H]UCB-J autoradiography in the frontal, temporal/parietal, occipital cortices and the hippocampus of WT and TG mice at 6, 12, and 18 months of age (Fig. 6A,B). Binding was age-dependently increased in 18-month-old TG mice compared to most other groups examined (Fig. 6A). In the temporal/parietal and occipital cortices, [^3^H]UCB-J levels were significantly higher in 18-month-old TG mice relative to age-matched WT controls (*P*<0.01) and to 6- and 12-month-old TG mice (*P*<0.05–0.001; LSD *post-hoc* tests). In the hippocampus, elevated binding levels were observed in 18- vs. 12-month-old TG animals (*P*<0.05). No differences were observed in the frontal cortex. Representative autoradiograms of [^3^H]UCB-J binding are shown in Fig. 6B. Factorial ANOVA revealed significant main effects of age [F(2,203)=11.5, *P*<0.001], genotype [F(1,203)=4.9, *P*<0.05] and brain region [F(3,203)=38.6, *P*<0.001] on the binding levels of [^3^H]UCB-J, along with significant genotype x age [F(2,203)=5.6, *P*<0.01] and genotype x age x region interaction effects [F(6,203)=2.3, *P*<0.05].

**Figure 6.**
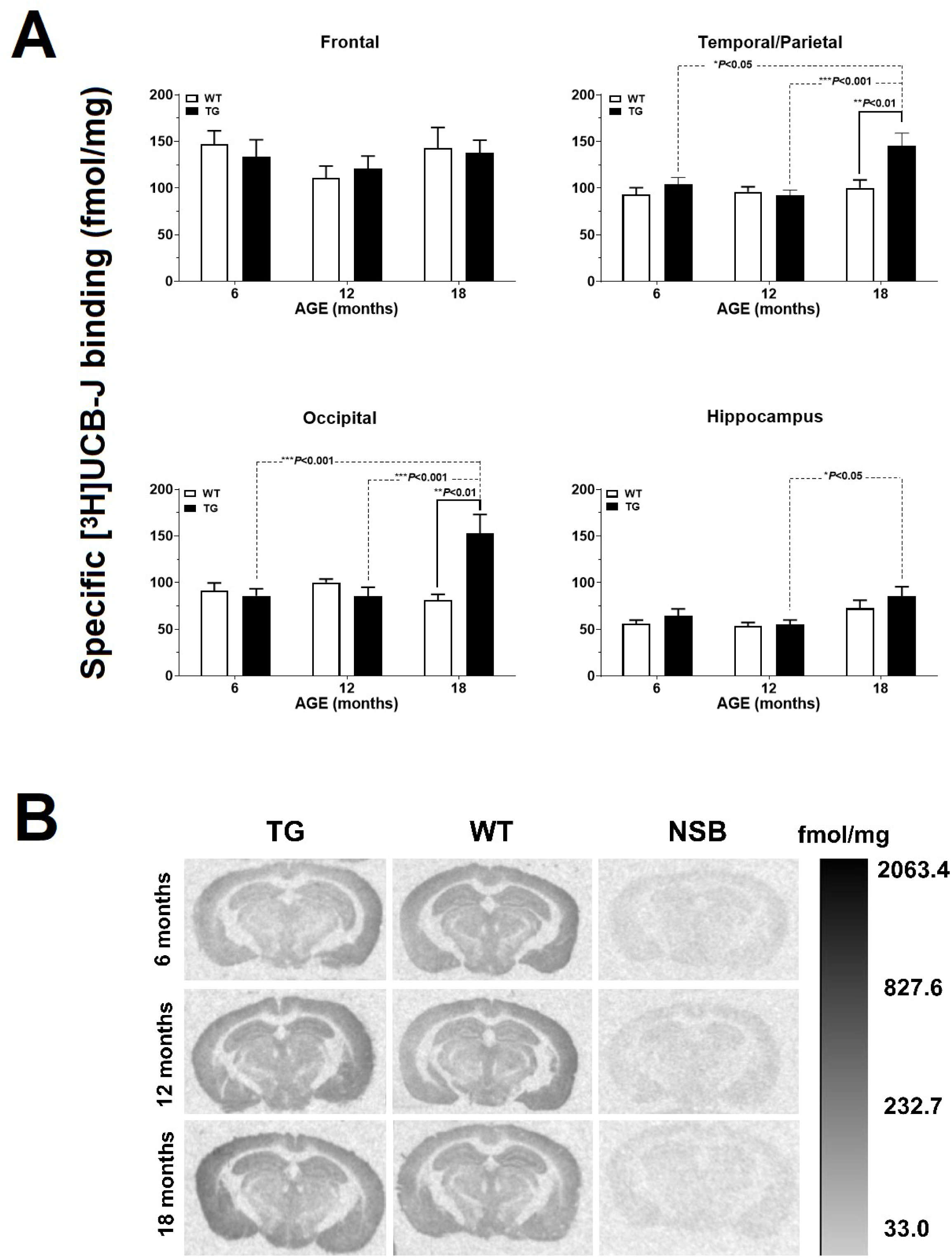
Age- and region-dependent increase in [^3^H]UCB-J binding in APP/PS1 mice. (A) Specific binding of [^3^H]UCB-J, a marker of presynaptic vesicle glycoprotein 2A (SV2A), was quantified by autoradiography in the frontal, temporal/parietal, occipital cortices, and hippocampus of wild-type (WT, open bars) and APP/PS1 transgenic (TG, filled bars) mice aged 6, 12, and 18 months. [^3^H]UCB-J binding was increased in TG mice, particularly in the temporal/parietal (P<0.05–0.001), occipital (P<0.01–0.001), and hippocampal (P<0.05) regions at 18 months, compared with age-matched WT mice and younger TG animals. No significant differences were detected in the frontal cortex. (B) Representative autoradiograms of [^3^H]UCB-J in TG and WT mice. Non-specific binding (NSB) was determined in adjacent sections incubated with 500 μΜ levetiracetam. The scale bar is a grayscale representation of binding intensity, calibrated in fmol/mg tissue equivalent.

### Age-dependent increase in Aβ plaque load in TG mice

Aβ plaque load increased significantly with age in TG mice in all brain areas examined (Fig. 7). Compared to 6-month-old TG mice, 12-month-old animals showed higher Aβ burden in the frontal and parietal/temporal cortices (*P*<0.001), the occipital cortex and the hippocampus (*P*<0.01; LSD *post-hoc* tests). A further increase in plaque load was observed at 18 months of age in all regions examined (*P*<0.001 for frontal, parietal/temporal, occipital; *P*<0.01 for hippocampus; LSD *post-hoc* tests). Among brain regions, the frontal cortex exhibited the highest plaque load across ages. Factorial ANOVA confirmed significant main effects of age [F(2,103)=153.8, *P*<0.001] and brain region [F(3,103)=10.1, *P*<0.001], with significant age x region interaction effects [F(6,103)=2.6, *P*<0.05].

**Figure 7.**
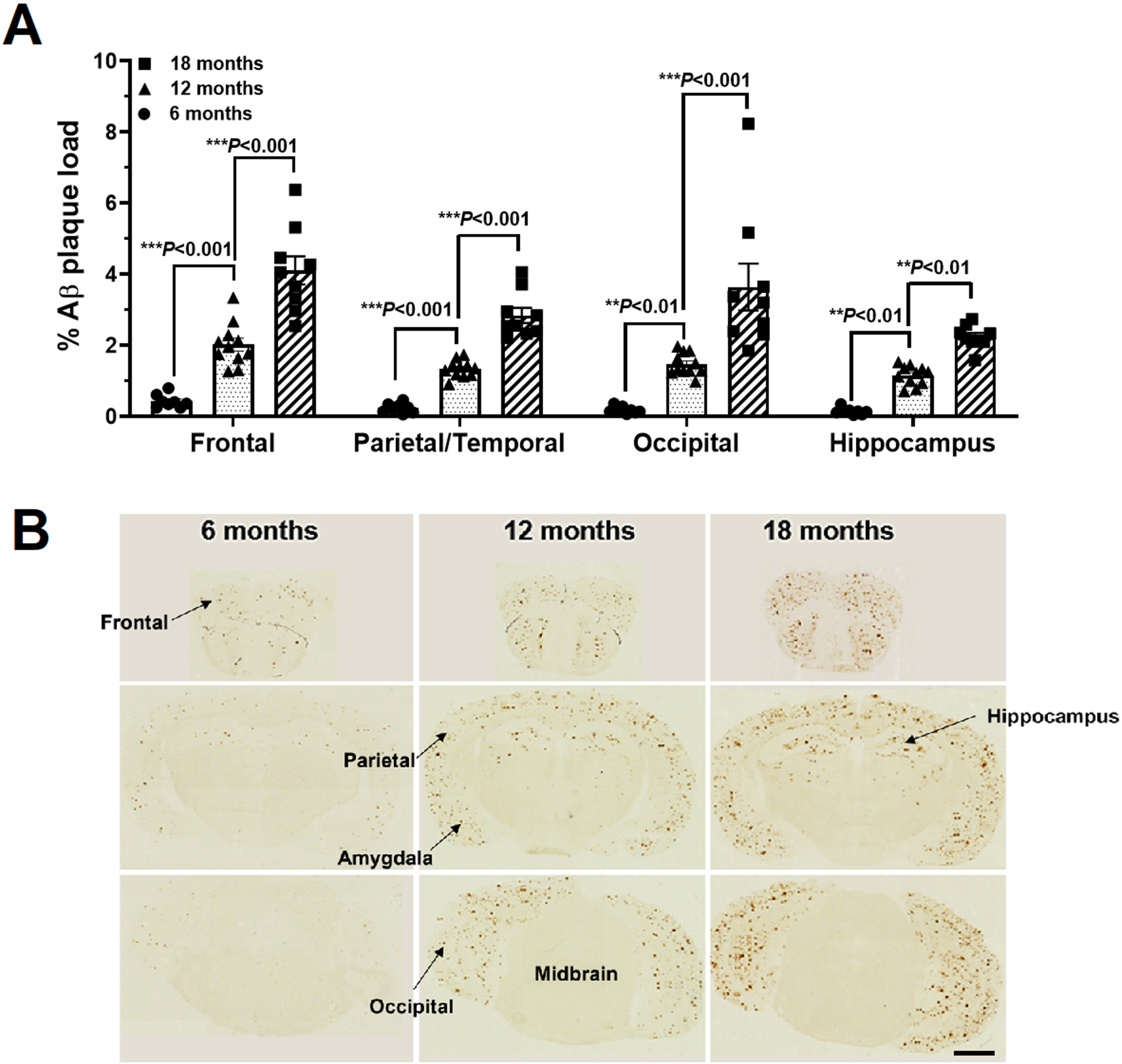
Age-dependent increase in Aβ plaque load in APP/PS1 mice. (A) Quantification of Aβ plaque load (% area covered by Aβ immunoreactivity) in the frontal, parietal/temporal, occipital cortices, and hippocampus of APP/PS1 transgenic mice aged 6, 12, and 18 months. Aβ deposition increased with age in all regions examined. (B) Representative immunohistochemical images showing progressive accumulation of Aβ plaques across brain regions in APP/PS1 mice at 6, 12, and 18 months of age. Labels indicate the anatomical regions analyzed. Scale bar = 2 mm.

### Associations between cognitive deficits and pathological markers in TG mice

Correlation analysis was performed in TG mice to examine the relationship between deficits in Barnes maze measures on day 11, including target block time, primary latency, and primary errors, with pathological markers of tau, Aβ, and synaptic density. Significant negative correlations were observed between target block time and Aβ plaque load (Fig. 8A) and [^18^F]flortaucipir binding (Fig. 8B) across cortical and hippocampal regions (r values = −0.43 to −0.63, *P*<0.05-0.001). [^3^H]UCB-J binding showed weaker associations, reaching significance in the frontal (r=-0.45, *P*<0.05) and parietal/temporal cortices (r=-0.38, *P*<0.05). There were no significant correlations between primary latency or primary errors and the examined pathological markers.

**Figure 8.**
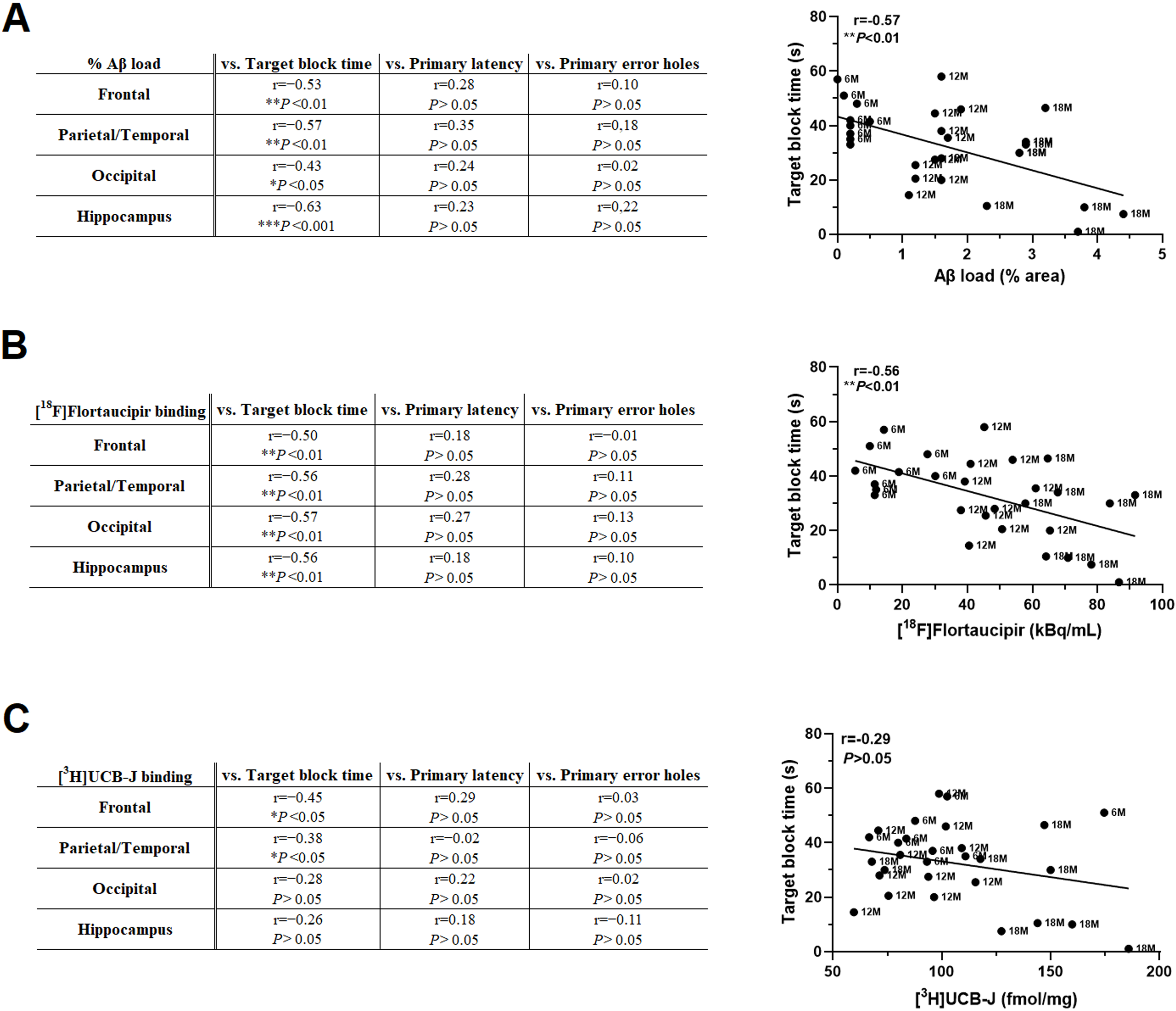
Correlations between markers of long-term memory retention and AD-like pathology in APP/PS1 mice. Pearson correlation analyses between Barnes maze performance and neuropathological measures in 6-, 12-, and 18-month-old APP/PS1 transgenic mice. Significant negative correlations were observed between Aβ plaque load and target block time in all cortical regions and the hippocampus (r = −0.43 to −0.63, *P*<0.05–0.001), indicating that higher amyloid burden was associated with poorer spatial memory performance. (B) [^18^F]Flortaucipir binding also correlated negatively with target block time in all regions analyzed (r = −0.50 to −0.57, *P*<0.01), linking increased tau accumulation with cognitive impairment. (C) [^3^H]UCB-J binding, a marker of synaptic density, showed weaker correlations, reaching significance only in the frontal (r = −0.45, *P*<0.05) and parietal/temporal cortices (r = −0.38, P<0.05). Scatter plots on the right illustrate representative correlations between target block time and the total levels of each pathological marker in the brain.

To account for the potential confounding effect of age, we next applied linear mixed-effects regression models to assess whether Aβ, tau and presynaptic density predicted target block time in TG mice. In cortical regions, Barnes maze performance was not significantly predicted by Aβ [F(1,13)=2.47, *P*=0.140], tau [F(1,13)=1.01, *P*=0.334] or [^3^H]UCB-J binding alone [F(1,13)=0.07, *P*=0.795], although the main effect of age approached significance [F(2,13)=3.69, *P*=0.054]. Importantly, the cortical model revealed two important interactions. First, a robust Aβ × tau effect [F(1,13)=8.02, *P*=0.014; estimate=-1.64], indicating that deficits in target block time were specifically associated with the combined presence of both pathologies in the cortex. Second, a significant tau × age interaction effect [F(2,13)=4.26, *P*<0.05], whereby tau binding was associated with increased target block time in 6-month-old TG mice (estimate=+4.51, *P*<0.05), an effect that reversed with advancing age, such that higher [^18^F]Flortaucipir binding predicted poorer performance at 18-month-old animals (estimate=-2.26, *P*<0.05). No significant effects were found for Aβ × age (*P*=0.216), UCB-J x age (*P*=0.599), or other higher-order interactions. In the hippocampus, target block time was not significantly predicted by age [F(2,13)=0.89, *P*=0.433], tau [F(1,13)= 0.00, *P*=0.997], Aβ [F(1,13)=0.14, *P*=0.711], or [^3^H]UCB-J binding [F(1,13)=0.02, *P*=0.880], and no significant interactions were detected (all *P* > 0.20).

## Discussion

The present study demonstrates that age-dependent impairments in spatial learning and memory in a mouse model of familial AD are most strongly associated with the combined accumulation of cortical Aβ and tau pathology. The pattern of deficits observed in APP/PS1 mice, with no impairment in the Barnes maze at 6 months, selective long-term memory impairment at 12 months, and profound learning and memory impairment at 18 months, mirrors the gradual progression of cognitive decline observed in AD patients [24, 25]. These results indicate that early disruptions in memory consolidation progress into broader learning deficits as AD pathology advances, reinforcing the close association between neuropathological burden and cognitive decline.

Deficits in learning and memory in APP/PS1 mice have been reported previously using the Barnes maze test [26–31]. However, differences in methodological design, including variations in the age of experimental animals, the number of task training days and trials, the lighting conditions and the use of proximal or distal cues, complicate direct comparisons between studies. In line with studies using the standard, hidden-target version of the Barnes maze, we show that learning and memory decline with age and increasing pathology in APP/PS1 mice [28, 29]. In particular, we were able to distinguish an early deficit in the long-term memory retention of 12-month-old TG mice from a later impairment in task acquisition at 18 months, indicating that memory consolidation declines before the capacity for learning a task is affected. This pattern mimics clinical findings across the AD continuum, where impairments in memory consolidation emerge early, preceding deficits in learning and broader cognition [32–34]. Interestingly, we also observed a mild but measurable effect of age on the long-term memory retention of 18-month-old WT animals (Fig. 2A & Fig. 3C), indicating that physiological ageing contributes to reduced cognitive performance, independent of Αβ and tau pathology. Therefore, our results show that the Barnes maze can distinguish physiological from pathological ageing, by detecting both the mild decline in spatial performance associated with ageing and the progressive impairment of learning and memory characteristic of AD. Importantly, the observed deficits are likely to reflect genuine cognitive dysfunction rather than sensory or gross motor limitations [35–38], since performance in the light/dark exploration test was normal and the number of holes visited during the probe trials did not differ between genotypes.

The mechanisms underlying progressive cognitive impairment in dominantly-inherited AD are associated with the accumulation of both plaques and tangles in the brain [39]. In line with this, we observed a progressive build-up of Aβ plaques in the cortex and the hippocampus of APP/PS1 mice, which paralleled the worsening of their cognitive performance. Similarly, [^18^F]flortaucipir binding revealed an age-dependent increase in tau signal exclusively in TG animals. Correlation analyses showed that higher Aβ plaque load and [^18^F]flortaucipir binding levels were each associated with worse long-term memory retention in the Barnes maze, while linear mixed-effects models revealed a significant interaction between Aβ and tau, even after controlling for age. These findings indicate that memory impairment in APP/PS1 mice is best explained by the synergistic effects of Aβ and tau on behavior, a pattern consistent with human PET imaging studies, showing that cognitive decline accelerates when tau pathology emerges in an amyloid-positive context [40–42].

The detection of tau-related effects in an amyloidosis model is noteworthy, as APP/PS1 mice are not typically considered to develop overt tauopathy. Nevertheless, recent evidence supports the notion that amyloid is sufficient to drive alterations in tau, especially with ageing. For example, we previously demonstrated that ageing and amyloidosis induce molecular and pathological tau changes in 18-24-month-old female APP/PS1 mice, comprising Gallyas-positive tau inclusions, correlations between argyrophilia and [^18^F]flortaucipir binding, and sarkosyl-insoluble tau hyperphosphorylation [11]. More recently, a meta-analysis of multiple AD mouse models, including APP/PS1 mice, showed that soluble tau phosphorylation levels are significantly increased in transgenic animals compared to controls, and that ageing moderates the magnitude of hyperphosphorylation, particularly within the proline-rich domain of tau in the neocortex [13]. Taken together with the present data, these results support a model in which the progressive accumulation of Aβ promotes tau pathology even in “amyloidosis-only” models, with tau burden interacting with Aβ and age to impair cognition. The significant tau × Aβ and tau × age interactions identified here suggest that both Aβ and tau pathologies should be targeted to halt the progression of AD.

Synaptic loss is a hallmark of AD pathology and one of the strongest correlates of cognitive decline in AD patients [43, 44]. In this study, synaptic density was quantified by [^3^H]UCB-J autoradiography for the presynaptic SV2A and showed region- and age-dependent increases in the temporal/parietal and occipital cortices of 18-month-old APP/PS1 mice compared to WT animals, whereas no genotype-induced effects were observed in the frontal cortex and the hippocampus. These observations contrast with *in vivo* SV2A PET imaging studies in humans, which consistently show reduced cortical and hippocampal uptake of [^11^C]UCB-J in AD [44–47]. However, autoradiographic analyses of hippocampal and cortical brain sections from sporadic AD patients reveal approximately 1.4-2.4-fold higher [^3^H]UCB-J binding than in age-matched controls [48], a finding that is consistent with the present data. The apparent discrepancy between the *in vivo* PET and tissue autoradiography results may be explained by factors beyond synaptic density, including *ApoE4* status [49, 50], altered UCB-J tissue accessibility, SV2A upregulation/reduced recycling, and off-target binding to phosphorylated tau species in subcellular fractions [48]. We have previously reported a negative correlation between AT8-positive tau immunoreactivity and [^3^H]UCB-J binding in human AD tissue, suggesting that SV2A autoradiography may partly track tau-associated changes [19]. [^3^H]UCB-J autoradiography may therefore not represent a direct measure of synaptic density, which could explain the weak associations observed between [³H]UCB-J signal and behavioral performance in the present study.

In addition to the discrepancies between UCB-J PET and autoradiography, several limitations should be acknowledged in our study. First, the behavioral assessment was limited to the hidden-target version of the Barnes maze test, which may not capture subtler early impairments that could emerge under different conditions or cue configurations. Second, while visual and motor function were controlled, factors such as motivation or stress sensitivity were not quantified and could influence performance, particularly in aged mice. Third, although strong correlations were observed between behavioral deficits and pathological markers, causality cannot be inferred, and contributions from other processes such as neuroinflammation or vascular dysfunction cannot be excluded. Finally, this study included only male mice, whereas our previous characterization of tau pathology was conducted in female TG animals [11], and so potential sex-dependent differences in the behavioral consequences of tau pathology remain to be investigated.

Despite these limitations, our combined behavioral, autoradiographic and histological data provide a robust framework for understanding the progression of Alzheimer-like pathology and its cognitive consequences in the APP/PS1 model of familial AD. Our findings demonstrate that cognitive decline in TG mice is closely linked to the interaction between amyloid and tau, highlighting the contribution of spontaneous, age-related tau aggregation within an amyloid context, and providing a relevant platform for evaluating disease-modifying interventions across distinct stages of disease progression.

## Acknowledgements

This study was funded by the University of Southern Denmark (SDU2020, CoPING AD: Collaborative Project on the Interaction between Neurons and Glia in AD).

## Reference list

1. Knopman, D.S., et al., Alzheimer disease. Nat Rev Dis Primers, 2021. 7(1): p. 33.

2. Pereira, J.B., et al., Untangling the association of amyloid-beta and tau with synaptic and axonal loss in Alzheimer’s disease. Brain, 2021. 144(1): p. 310–324.

3. Terry, R.D., et al., Physical basis of cognitive alterations in Alzheimer’s disease: synapse loss is the major correlate of cognitive impairment. Ann Neurol, 1991. 30(4): p. 572–80.

4. Arnsten, A.F.T., et al., An integrated view of the relationships between amyloid, tau, and inflammatory pathophysiology in Alzheimer’s disease. Alzheimers Dement, 2025. 21(8): p. e70404.

5. Gallego-Rudolf, J., et al., Synergistic association of Abeta and tau pathology with cortical neurophysiology and cognitive decline in asymptomatic older adults. Nat Neurosci, 2024. 27(11): p. 2130–2137.

6. He, Z., et al., Amyloid-beta plaques enhance Alzheimer’s brain tau-seeded pathologies by facilitating neuritic plaque tau aggregation. Nat Med, 2018. 24(1): p. 29–38.

7. Kadamangudi, S., et al., Amyloid-beta oligomers increase the binding and internalization of tau oligomers in human synapses. Acta Neuropathol, 2024. 149(1): p. 2.

8. Simoes-Pires, E.N., et al., Synergistic effects of the Abeta/fibrinogen complex on synaptotoxicity, neuroinflammation, and blood-brain barrier damage in Alzheimer’s disease models. Alzheimers Dement, 2025. 21(5): p. e70119.

9. Kim, B.H., et al., Second-generation anti-amyloid monoclonal antibodies for Alzheimer’s disease: current landscape and future perspectives. Transl Neurodegener, 2025. 14(1): p. 6.

10. Zhong, M.Z., et al., Updates on mouse models of Alzheimer’s disease. Mol Neurodegener, 2024. 19(1): p. 23.

11. Metaxas, A., et al., Ageing and amyloidosis underlie the molecular and pathological alterations of tau in a mouse model of familial Alzheimer’s disease. Sci Rep, 2019. 9(1): p. 15758.

12. Babcock, A.A., et al., Cytokine-producing microglia have an altered beta-amyloid load in aged APP/PS1 Tg mice. Brain Behav Immun, 2015. 48: p. 86–101.

13. Kourti, M. and A. Metaxas, A systematic review and meta-analysis of tau phosphorylation in mouse models of familial Alzheimer’s disease. Neurobiol Dis, 2024. 192: p. 106427.

14. Barnes, C.A., Memory deficits associated with senescence: a neurophysiological and behavioral study in the rat. J Comp Physiol Psychol, 1979. 93(1): p. 74–104.

15. Jankowsky, J.L., et al., Co-expression of multiple transgenes in mouse CNS: a comparison of strategies. Biomol Eng, 2001. 17(6): p. 157–65.

16. Mygind, L., et al., Tumor Necrosis Factor (TNF) Is Required for Spatial Learning and Memory in Male Mice under Physiological, but Not Immune-Challenged Conditions. Cells, 2021. 10(3).

17. Thompson, S., et al., Light aversion in mice depends on nonimage-forming irradiance detection. Behav Neurosci, 2010. 124(6): p. 821–7.

18. Shoup, T.M., et al., A concise radiosynthesis of the tau radiopharmaceutical, [(18) F]T807. J Labelled Comp Radiopharm, 2013. 56(14): p. 736–40.

19. Metaxas, A., et al., Increased Inflammation and Unchanged Density of Synaptic Vesicle Glycoprotein 2A (SV2A) in the Postmortem Frontal Cortex of Alzheimer’s Disease Patients. Front Cell Neurosci, 2019. 13: p. 538.

20. Lynch, B.A., et al., The synaptic vesicle protein SV2A is the binding site for the antiepileptic drug levetiracetam. Proc Natl Acad Sci U S A, 2004. 101(26): p. 9861–6.

21. Edison, P., et al., Amyloid, hypometabolism, and cognition in Alzheimer disease: an [11C]PIB and [18F]FDG PET study. Neurology, 2007. 68(7): p. 501–8.

22. Chen, B., et al., PET Imaging in Animal Models of Alzheimer’s Disease. Front Neurosci, 2022. 16: p. 872509.

23. Snellman, A., et al., Longitudinal amyloid imaging in mouse brain with 11C-PIB: comparison of APP23, Tg2576, and APPswe-PS1dE9 mouse models of Alzheimer disease. J Nucl Med, 2013. 54(8): p. 1434–41.

24. Koscik, R.L., et al., Amyloid duration is associated with preclinical cognitive decline and tau PET. Alzheimers Dement (Amst), 2020. 12(1): p. e12007.

25. Payton, N.M., et al., Trajectories of cognitive decline and dementia development: A 12-year longitudinal study. Alzheimer’s & Dementia, 2023. 19(3): p. 857–867.

26. Hulshof, L.A., et al., Both male and female APPswe/PSEN1dE9 mice are impaired in spatial memory and cognitive flexibility at 9 months of age. Neurobiol Aging, 2022. 113: p. 28–38.

27. Lesuis, S.L., et al., Targeting glucocorticoid receptors prevents the effects of early life stress on amyloid pathology and cognitive performance in APP/PS1 mice. Transl Psychiatry, 2018. 8(1): p. 53.

28. O’Leary, T.P. and R.E. Brown, Visuo-spatial learning and memory deficits on the Barnes maze in the 16-month-old APPswe/PS1dE9 mouse model of Alzheimer’s disease. Behav Brain Res, 2009. 201(1): p. 120–7.

29. Reiserer, R.S., et al., Impaired spatial learning in the APPSwe + PSEN1DeltaE9 bigenic mouse model of Alzheimer’s disease. Genes Brain Behav, 2007. 6(1): p. 54–65.

30. Xu, L., et al., Deficits in N-Methyl-D-Aspartate Receptor Function and Synaptic Plasticity in Hippocampal CA1 in APP/PS1 Mouse Model of Alzheimer’s Disease. Front Aging Neurosci, 2021. 13: p. 772980.

31. Zhao, Y., et al., Spatial Training Attenuates Long-Term Alzheimer’s Disease-Related Pathogenic Processes in APP/PS1 Mice. J Alzheimers Dis, 2022. 85(4): p. 1453–1466.

32. Johnson, D.K., et al., Longitudinal study of the transition from healthy aging to Alzheimer disease. Arch Neurol, 2009. 66(10): p. 1254–9.

33. O’Connor, A., et al., Quantitative detection and staging of presymptomatic cognitive decline in familial Alzheimer’s disease: a retrospective cohort analysis. Alzheimers Res Ther, 2020. 12(1): p. 126.

34. Ohno, M., Accelerated long-term forgetting: A sensitive paradigm for detecting subtle cognitive impairment and evaluating BACE1 inhibitor efficacy in preclinical Alzheimer’s disease. Front Dement, 2023. 2: p. 1161875.

35. Huang, H., et al., Isolation Housing Exacerbates Alzheimer’s Disease-Like Pathophysiology in Aged APP/PS1 Mice. Int J Neuropsychopharmacol, 2015. 18(7): p. pyu116.

36. Savonenko, A., et al., Episodic-like memory deficits in the APPswe/PS1dE9 mouse model of Alzheimer’s disease: relationships to beta-amyloid deposition and neurotransmitter abnormalities. Neurobiol Dis, 2005. 18(3): p. 602–17.

37. Sood, A., et al., The effects of JWB1-84-1 on memory-related task performance by amyloid Abeta transgenic mice and by young and aged monkeys. Neuropharmacology, 2007. 53(5): p. 588–600.

38. Wiesmann, M., et al., Hypertension, cerebrovascular impairment, and cognitive decline in aged AβPP/PS1 mice. Theranostics, 2017. 7(5): p. 1277–1289.

39. Bateman, R.J., et al., Clinical and biomarker changes in dominantly inherited Alzheimer’s disease. N Engl J Med, 2012. 367(9): p. 795–804.

40. Cody, K.A., et al., Characterizing brain tau and cognitive decline along the amyloid timeline in Alzheimer’s disease. Brain, 2024. 147(6): p. 2144–2157.

41. Ossenkoppele, R., et al., Tau PET positivity in individuals with and without cognitive impairment varies with age, amyloid-beta status, APOE genotype and sex. Nat Neurosci, 2025. 28(8): p. 1610–1621.

42. Ossenkoppele, R., et al., Amyloid and tau PET-positive cognitively unimpaired individuals are at high risk for future cognitive decline. Nat Med, 2022. 28(11): p. 2381–2387.

43. DeKosky, S.T. and S.W. Scheff, Synapse loss in frontal cortex biopsies in Alzheimer’s disease: correlation with cognitive severity. Ann Neurol, 1990. 27(5): p. 457–64.

44. Mecca, A.P., et al., Synaptic density and cognitive performance in Alzheimer’s disease: A PET imaging study with [(11) C]UCB-J. Alzheimers Dement, 2022. 18(12): p. 2527–2536.

45. Chen, M.K., et al., Assessing Synaptic Density in Alzheimer Disease With Synaptic Vesicle Glycoprotein 2A Positron Emission Tomographic Imaging. JAMA Neurol, 2018. 75(10): p. 1215–1224.

46. Mecca, A.P., et al., In vivo measurement of widespread synaptic loss in Alzheimer’s disease with SV2A PET. Alzheimers Dement, 2020. 16(7): p. 974–982.

47. Nilsson, J., et al., Associations between fluid biomarkers and PET imaging ([11C]UCB-J) of synaptic pathology in Alzheimer’s disease. Alzheimers Dement, 2025. 21(7): p. e70403.

48. Kumar, A., M. Scarpa, and A. Nordberg, Tracing synaptic loss in Alzheimer’s brain with SV2A PET-tracer UCB-J. Alzheimers Dement, 2024. 20(4): p. 2589–2605.

49. Mikkelsen, J.D., et al., Synaptic vesicle glycoprotein 2A (SV2A) levels in the cerebral cortex in patients with Alzheimer’s disease: a radioligand binding study in postmortem brains. Neurobiol Aging, 2023. 129: p. 50–57.

50. Mikkelsen, J.D., P. Linde-Atkins, and B.A. Pazarlar, Higher level of [(3)H]UCB-J binding in ApoE E4 allele carriers with Alzheimer disease. Neurosci Lett, 2025. 849: p. 138135.

