## Supplementary for "Age-specific Aβ-Tau interactions underlie spatial memory deficits in an APP/PS1 amyloidosis model"

**Supplementary Information**

*[^11^C]PIB autoradiography*

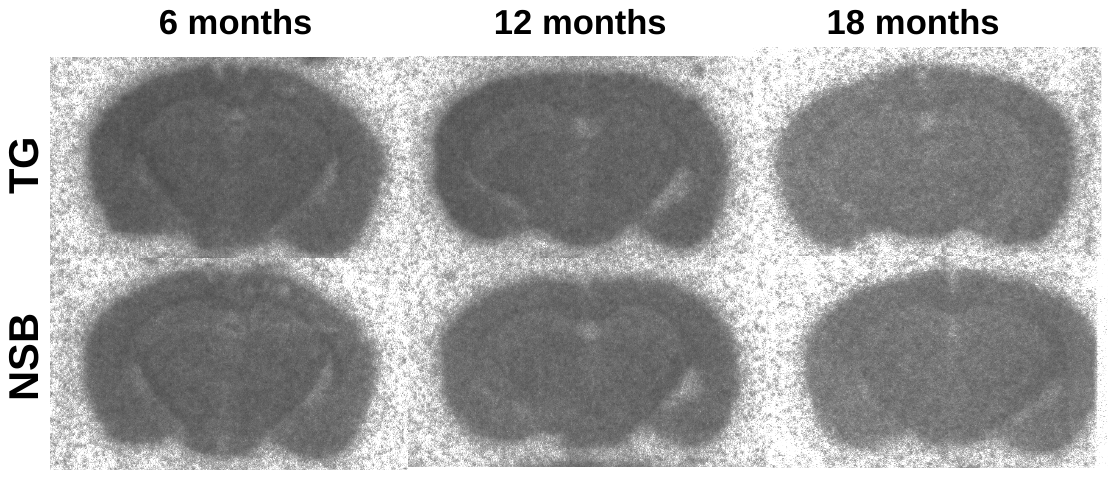
Brain sections were thawed to RT and incubated for 40 min with [^11^C]PIB (636 kBq/mL, specific activity: 360 GBq/μmol), in 50 mM Tris-buffered saline (TBS) containing 5% ethanol. Adjacent sections were incubated with identical amounts of radioligand in the presence of 10 μM non-radiolabelled PIB, to calculate NSB. Following rapid washing in ice-cold 50 mM TBS + 5% ethanol (1 min), TBS (2 min) and a dip in ice-cold dH_2_O. After rapid drying under a stream of cold air, the sections were placed in light-tight cassettes and exposed against FUJI multi-sensitive phosphor screens for 30 min (BAS-IP SR2025, GE Healthcare Life Sciences). Images were developed in a BAS-5000 phosphor-imager at 25 μm resolution. Values of specific binding were derived after subtraction of non-specific from total binding images.

**Supplementary Fig. 1**: Representative [^11^C]PIB autoradiograms in TG mice, showing lack of specific binding.

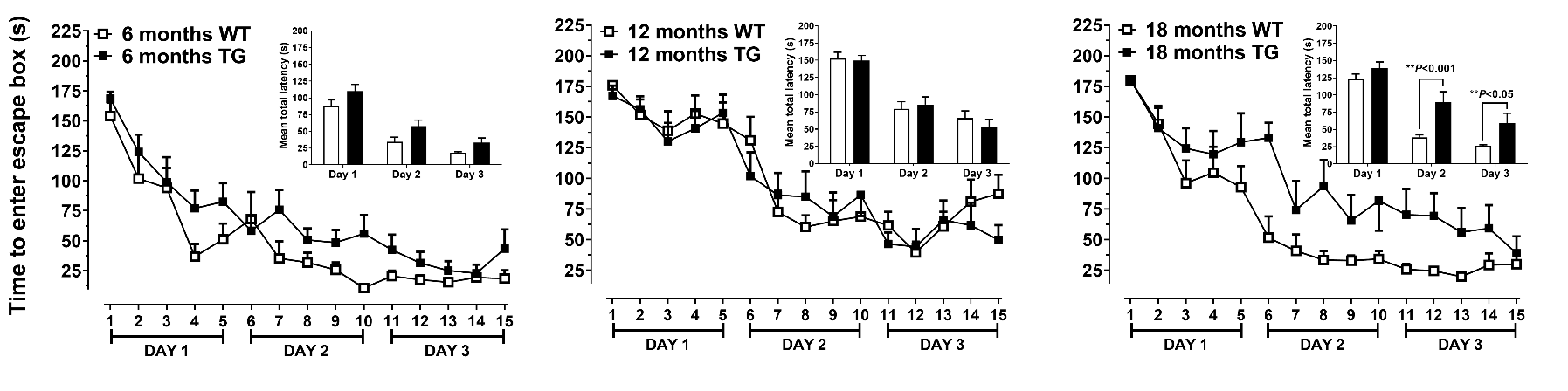
*Barnes Maze Acquisition Phase: Total Latency*

**Supplementary Fig. 2**: Mean latency to enter the escape box across 15 trials of the Barnes maze training in 6-, 12-, and 18-month-old WT and APP/PS1 (TG) mice. Insets show average total latency per training day. APP/PS1 mice exhibited delayed task acquisition at 18 months (*P*<0.001–0.05 vs. WT).
